# ProteomeScout v4: An open-source, harmonized, ecosystem for post-translational modifications

**DOI:** 10.64898/2026.10.01.756115

**Authors:** Sam Crowl, Saqib Rizvi, Logan Campbell, Alekhya Kandoor, Kristen M. Naegle

## Abstract

**Motivation:** There are many databases of post-translational modifications, but each has different protein and PTM coverage, making the current state of post-translational modifications difficult to obtain, assess, or harmonize. ProteomeScout has been a relational database that harmonized a myriad of resources to address this challenge. In this update, we wished to create a lightweight, sustainable open source resource for the community that retained the power of harmonization, protein visualization, and Python tools for interacting with the database.

**Results:** ProteomeScout-v4 presents an ecosystem of open-source tools for aggregating and updating proteins and post-translational modifications and presenting the web-based and code-based access tools for interacting with the data. ProteomeScout has moved from using a complex relational database to a flat-text file format. The database file update is version controlled on FigShare and all features have processes for continuous updating, including migrating PTMs to updated protein records, when external databases are updated. The open-source nature and flexibility allow for user-based control and community-based feature integration. Fast and easy analysis is possible with the API, with example analyses provided here.

**Availability and implementation:** ProteomeScout-v4 web interface is available at https://proteomescout.research.virginia.edu/; the web interface package is available at: https://github.com/NaegleLab/proteomescout-v4; the API is available at https://github.com/NaegleLab/ProteomeScoutAPI; database update code is available at https://github.com/NaegleLab/ProteomeScout-v4_Update; and current database files are maintained at: https://doi.org/10.6084/m9.figshare.31129045.

## Introduction

Post-translational modifications (PTMs), the covalent addition of chemical groups to proteins, can profoundly influence protein function, stability, localization, and interactions. Providing regulatory information that is not encoded directly by the central dogma of biology, PTMs are essential for: directing proteins to their appropriate cellular destinations (e.g. glycosylation for cell-surface presentation or fatty acid additions, like palmitoylation, for membrane anchoring); mediating specific protein-protein interactions (e.g. acetylation, methylation, and phosphorylation); and regulating protein lifetime (e.g. SUMOylation and ubiquitination). The advent of tools for su!cient PTM enrichment and measurement by mass spectrometry-based approaches led to a massive expansion of our understanding of the extent of PTMs within the human proteome (8), as well as in other model organisms, such as mouse, yeast, and fly. Mass spectrometry-based profiling of PTMs is now so well established that it is regularly part of the molecular multiomic profiling of cells and tissues, including broad-scale efforts in cancer (14).

PTMs and various information regarding PTMs and their proteins, and each has distinct approaches for providing updates. Some early databases, which are still important today, relied heavily on hand curation and are no longer updated (e.g. phosphoELM (6) and HPRD (7)). Some databases, such as UniProt (15) and PhoshpoSitePlus (9) are more continuously updated (though at the time of this article PhosphoSitePlus has suspended academic licenses). There are databases that seek to integrate multiple PTM resources, such as dbPTM, which is updated by the publication of a major revision every few years (2). Due to their different emphasis on collecting PTM information, no single database captures all annotated PTMs (Fig. 1) and integrating across resources is non-trivial problem – each resource is referenced differently (e.g. different versions of the same protein database, or even different protein databases). At its inception, we developed PTMScout (13) for the purpose of harmonizing PTM information from databases and experiments, along with contextualizing those PTMs within proteins, which was re-released as an update with expanded tools for protein visualization (12), along with an API for interacting with a flat-text version of the database (8). However, a complex, relational database, with web-based interaction tools, creates issues for long-term maintenance in the academic setting. We have re-envisioned ProteomeScout as an open-source set of tools that works entirely on the flat-text format. In this open-source, simple format of a PTM database, rapid development by the community, regular updates, and database/feature personalization are now easily obtainable. As part of this v4 update, we migrated and updated protein data, incorporated new data features (exons (3) and SH2 domain predictions (11)), and added new tools and features to the web site and API (such as the fast annotation of PTM experiments, dataset preparation and plotting for KSTAR (5; 4), and resizing and export of protein track images.

**Figure 1.**
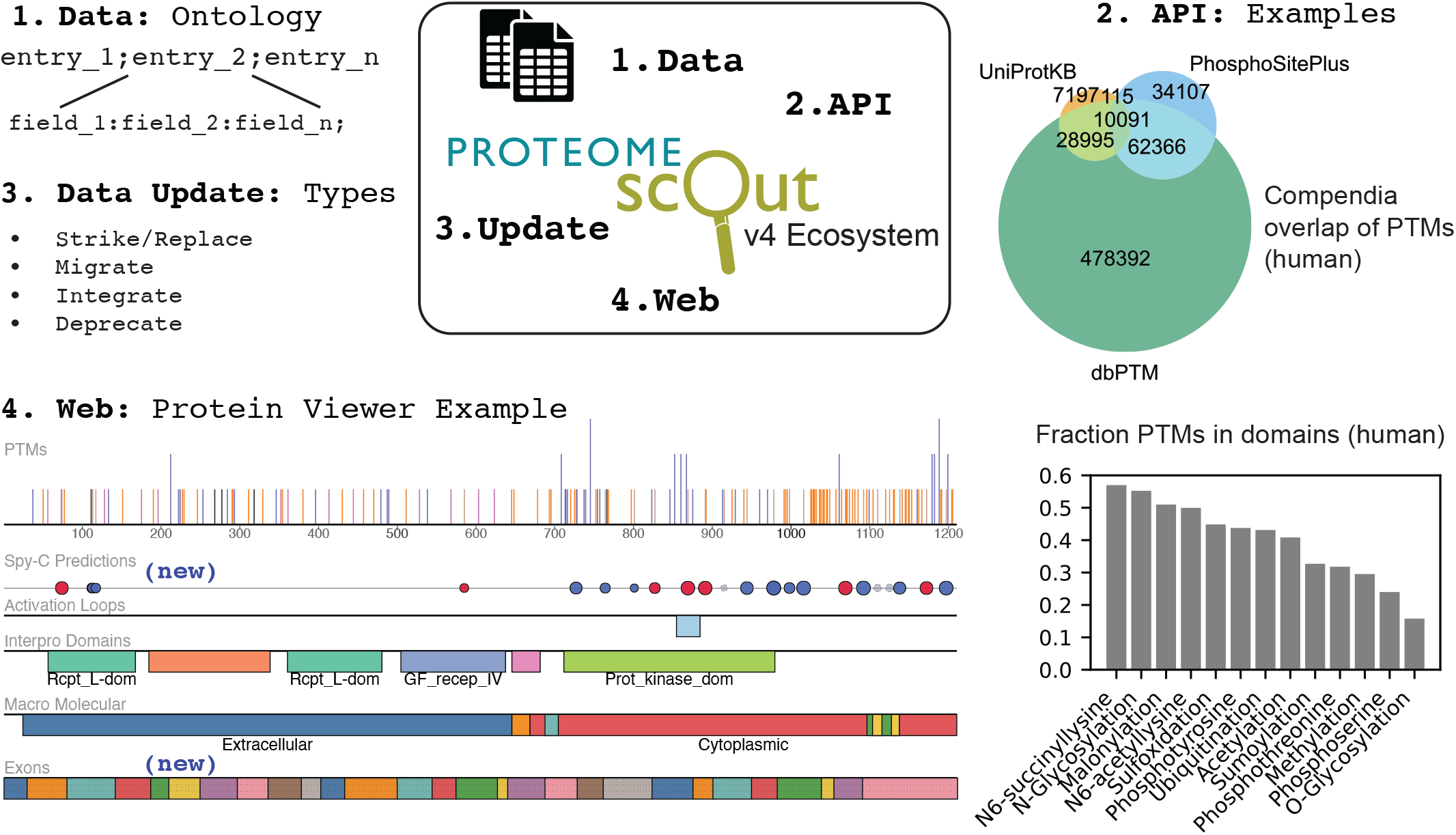
ProteomeScout-v4 Ecosystem. An overview of the ecosystem that consists of a data.tsv file and a citations.tsv file, containing all ProteomeScout harmonized data, current features, and PTMs. The data file consists of one row per protein and feature columns with customized feature fields following colon separation for each entry and semicolon separated entries. When integrating a new data feature, the first integration is required to add the new feature to all available and applicable records under a defined ontology, which is then integrated as a method for further updating in the future. Depending on the feature, it is integrated into the API code for parsing and retrieval, and added to the annotation and species reference generation code in the API, which is integrated directly into the web-based interface. If applicable, code for parsing and visualizing the new feature is added for the protein viewer (web-based). Here, an example record for EGFR (P00533) demonstrates example feature information and tracks available under the Protein Viewer (tracks not pictured are UniProt domains, and secondary structure). PTMs are indicated, colored based on amino acid (tan: tyrosine; orange: serine/threonine; blue lysine; gray methionine). In addition to hiding tracks, PTMs can be filtered on the viewer and zoom, with SVG export. Additionally, ProteomeScout web interface provides non-programmatic access to two KSTAR (a tool for predicting kinase activity from phosphoproteomic datasets (5)) features – dataset preparation for using in KSTAR-Galaxy or Python KSTAR, and plotting of KSTAR results. Two analyses performed for the human proteome using the API are indicated, including: 1) the overlap of PTMs in three major compendia (UniProtKB (15), PhosphoSitePlus (9), and dbPTM (2) and 2) the fraction of PTMs located within domains for PTMs with more than 2,000 entries in the human proteome.

## Methods

### ProteomeScout ecosystem

ProteomeScout-v4 is an open source ecosystem, composed of four key components: 1) database flat text files (a harmonized database file and a citations file that associates PTM data with their sources), 2) a Python API – to interact with, and extract information from, the database files, 3) a light weight web interface that interacts with the database files for visualization and non-programmatic use of the database, and 4) code for database updates and integration. In this version of ProteomeScout, we performed a significant update, migrating prior versions of the database to current protein records and annotations and forming the basis for robust, long-term, updates of the records. This harmonization, with its ability to migrate records as the reference proteomes change, is a key asset to the field for long-term maintenance of PTM records. As an ecosystem, the open-source code and the controlled database formatting allow for community-based expansion, along with the use of personalized access (Fig. 1).

For example, a research lab could use the ecosystem to harmonize and track all data within the lab, running their own personalized web service internally, along with excluding versions of data in the public database that are not of interest.

### Data features of ProteomeScout

The database contains information from resources of two types:

1) protein-level information (Gene Ontology, sequence, accessions, domains, and structure elements) and 2) site-level information (PTMs and annotations or predictions related to PTMs). Protein information is centralized around UniProt records (15) and we have added database flags to indicate canonical records, allowing simple analysis of a species-level proteome. We capture the complete non-redundant proteomes for human and mouse, including records with no recorded PTMs – other species protein records are added as PTMs are associated with them. The ProteomeScout-v4 update modernized the types of protein-level information captured, including moving to InterPro as the primary source of domain information, based on code in the CoDIAC project (10) and unifying information by using UniProt as the primary resource of sequence, structure, and Gene Ontology terms. In this version, we integrated new features, including an updated kinase activation loop prediction approach (Supplementary Methods), exons based on ExonPTMapper (3), and predictions of SH2 domain interaction sites from SpY-C (11). For PTM databases that are regularly updated (PhosphoSitePlus (9) and UniProt (15)), or when other databases undergo new publication and a single update push, we integrate these updates into the database record. The process for new PTM data integration requires first formatting the incoming dataset into a standard format (UniProt accession and either a site or a modified peptide). During integration, if a new protein record is encountered, the protein is added to the database, along with associated PTMs and protein-relevant information. Otherwise, if it encounters a novel PTM, then that PTM is added to the protein record, or if the PTM exists already, the citation information is inserted into the existing record. We never remove information from the database record, but instead use a flag in the citations file that allows users to indicate a deprecated dataset. These citation flags are cross-referenced when generating protein viewers, or retrieving data from the database in the API – effectively removing it from analysis, unless overridden by an API request. Supplementary Data, as well as online documentation, fully detail data features, sources, ontologies that define how fields are stored, and the process for keeping this data updated.

As an open resource intended for community development, we created an ecosystem process for new feature integration (Fig. 1). The addition of a new database feature and its integration involves: defining the field information, adding the parsing of the field to the API, integrating the visualization of the information on the web resource, and adding the process for future updates to the update code. As an example, in this update we added exon details (a protein-level record) that will enhance analysis related to splicing and PTMs (3) and predictions from a new approach that classify the likelihood that tyrosine phosphorylation is recognized by by an SH2 domain (SpY-C) (11). We defined key information that was necessary to communicate these elements (e.g. the Ensembl Exon ID, boundaries, and a boolean field to indicate if the exon is constitutive), defined an ontology to store this information (exon id:start:end:constitutive), which is then integrated into the API for record parsing, as well as integration into key features (Dataset Annotation and Species Reference File generation, both API features that are integrated into the web resource). Finally, we integrated methods for updating this data in the database and incorporated a track-based visualization of these elements as part of the Protein Viewer on the web resource (Fig. 1). These steps define everything necessary to provide new protein or PTM-level information in a long-term sustainable manner with high transparency to users. Selection of information to encode for any new field is highly flexible, which enhances downstream use cases.

## Example Applications

### ProteomeScout web tools

The ProteomeScout website gives access to the database and directly integrates key API features. It includes: 1) protein visualization – a track-based approach with the ability to filter tracks and PTM types, includes zoom, and figure export (in scalar vector graphic format with width scaling), 2) dataset annotation – given any comma separated file containing at least a UniProt identifier and a site or a peptide with a PTM indicated, this tool will append all available feature information to that dataset, which includes information such as: whether and in what datasets that modification has been observed, protein features the modification falls within (domains, macromolecular, structure) and protein information (gene ontology, domains, etc.), and 3) dataset preparation – given a comma separated file, this tool will map non-UniProt identifiers and process common designations in PTM experiment files into a common format, such as used for kinase activity analysis and dataset integration into ProteomeScout. ProteomeScout additionally provides non-programmatic plotting tools for KSTAR activity analysis (5; 4). Finally, ProteomeScout downloads include the specific database files (also versioned on FigShare) and species-specific datasets that present the data in a PTM-centric approach, which allows for fast analysis of proteome-level information.

### ProteomeScout API

Working directly with the database file provides flexible and ample analysis opportunities across any feature or PTM within the file and the API provides a fast, Python-based interaction interface. Focused on returning record information, based on UniProt ID, a user can quickly evaluate information about individual or groups of protein records – such as iterating across all protein in a species. Figure 1 includes two examples of API-based full database and proteome-level analysis. Using the API and the latest version of ProteomeScout data (version 9), we analyzed the overlap between the largest and most recent compendia of PTMs for the human proteome (Fig. 1). This highlights the need for harmonization – each database is not fully captured by any other database and so no single resource allows for comprehensive analysis. The ability of ProteomeScout to harmonize records, along with continuous updates, is key to enabling robust analysis. The second example is an exploration of PTM-specific behavior, measuring the fraction of PTMs that fall within defined InterPro domains (1), for any PTM with more than 2,000 entries in the human proteome (Fig. 1). This highlights the important differences in behavior, even within similar PTM families. For example, tyrosine phosphorylation and N-linked glycosylation occur at a much higher rate in domains than their counterparts, hosphothreonine, phosphoserine, and O-linked glycosylation, which are much less likely to be located in domains.

## Conclusion

ProteomeScout is a comprehensive database of post-translational modifications, harmonizing the records across other major compendia. It uniquely provides an open-source environment with flexibility for adding new features, consistent updates, customization, and fast programmatic analysis. The website provides programming-free tools for annotating PTM datasets and preparing them for other downstream applications.

## Abbreviations

PTMI: post-translational modification
SVGI: scalar vector graphic
PSPI: PhosphoSitePlu

## Conflicts of interest

The authors declare that they have no competing interests.

## Funding

Research reported in this publication was supported by the National Institute Of General Medical Sciences of the National Institutes of Health under Award Number R35GM138127 and the National Cancer Institute under Award Number U01CA284193. The content is solely the responsibility of the authors and does not necessarily represent the official views of the National Institutes of Health.

## Data availability

The data underlying this article are available at Figshare at https://doi.org/10.6084/m9.figshare.31129045 and can be accessed with specific version numbers by physical download or by using the API at https://github.com/NaegleLab/ProteomeScoutAPI. The current data can be downloaded directly from the Downloads page at https://proteomescout.research.virginia.edu/.

## Author contributions statement

All authors made code and intellectual contributions. K.M.N wrote the manuscript and performed analysis.

## Notes

https://proteomescout.research.virginia.edu/

https://github.com/NaegleLab/ProteomeScoutAPI

https://doi.org/10.6084/m9.figshare.31129045

https://github.com/NaegleLab/proteomescout-v4

https://github.com/NaegleLab/ProteomeScout-v4_Update

### Competing Interest Statement

The authors have declared no competing interest.

