## Supplemental Methods for "ProteomeScout v4: An open-source, harmonized, ecosystem for post-translational modifications"

### Supplementary Methods for ProteomeScout-v4

#### Kinase Activation Loop Prediction

We created a reference fasta file of 1,050 kinase domains taken from across human, mouse, fly, and yeast species. We used mafft-linsi to align these, creating the global reference alignment. Kinase modification was used to change the size and composition to ensure that sufficient coverage of diversity allowed all human, mouse, rat, cow, fly, and yeast kinases to be aligned relative to the global reference. We then manually annotated the conserved APE and DFG flanking sequence positions in the reference alignment and kinase domains are extracted for protein records and aligned to this reference. We annotate the activation loop as having a start and end if APE and DFG are matched sufficiently to those reference position flanking sites. We annotate with high confidence label if we find both start and end flanking sequences with small degree of degeneracy. We label the activation loop as medium confidence if global alignment quality is good and at least one of the flanking sequences is matched. We return no activation loop if quality is poor or insufficient match to the tripeptide matches in the flanking sequences.

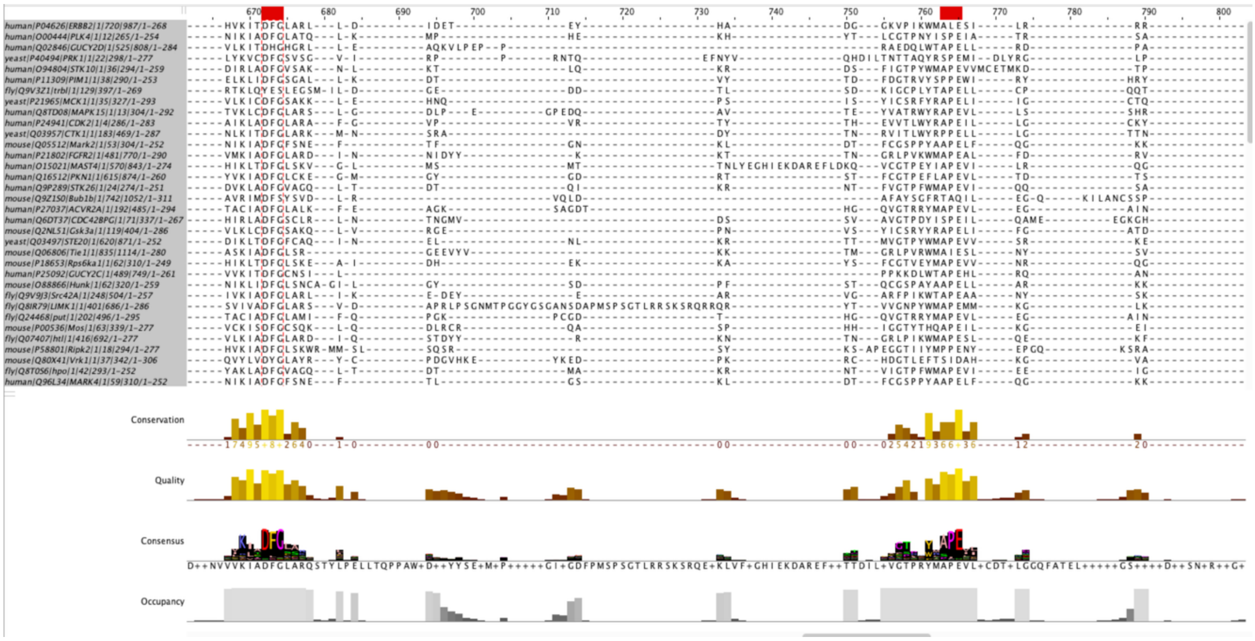


**Figure 1: Activation loop in reference alignment. DFG and APE conserved and highlighted in this reference alignment between 672 and 765.**
